# A nanobody-gold conjugate for the detection of GFP-tagged synaptic proteins by dual light-electron microscopy

**DOI:** 10.1101/2025.10.08.681152

**Authors:** Charles Ducrot, Cécile Lemoigne, Paul Lecompte Saint-Jean, Tiffany Cloâtre, Sophie Daburon, Kerstin Seiling, Leonie Mohrmann, Astrid Rohlmann, Markus Missler, Rémi Fronzes, Daniel Choquet, Matthieu Sainlos, Olivier Thoumine

## Abstract

To meet the constantly improving spatial resolution offered by advanced microscopy techniques to study sub-cellular structures in biology, there is a need for small, monovalent probes that label proteins of interest with high specificity and minimal distance to the target, and are compatible with various imaging modalities. In this direction, we designed a strategy to generate minimal-size probes composed of a controlled 1:1 conjugate between a small domain binder and a 1.4 nm-gold nanoparticle with direct access to fluorescent labelling for dual light-electron microscopy. Our approach was applied to the widely used single-domain antibody against GFP (GBP). The modified GBP-gold conjugate retained normal binding to purified GFP in vitro, specifically labelled COS-7 cells and neurons expressing GFP-tagged synaptic membrane proteins, and penetrated readily into tight cell-cell contacts including neuronal synapses. The optional fluorescence labelling with a second ALFA nanobody allowed dSTORM imaging, while the silver-enhanced nanogold particle detected in TEM was used to characterize the number and nanoscale organization of individual proteins in the synaptic cleft. We counted a small number of endogenous neurexins in the pre-synapse and a larger number of AMPA receptors in the post-synapse, often aligned in nanodomains. This GBP-gold probe thus emerges as a potent tool to label an ever-increasing repertoire of GFP-tagged proteins in numerous biological organisms and models.

## Introduction

Many biological functions are performed at the level of sub-micron cellular structures that are composed of highly organized multi-protein complexes. For example, in neuronal synapses, adhesion molecules and glutamate receptors form nanodomains precisely aligned with neurotransmitter release zones ^1–5^, an organization that is essential to synaptic transmission. To decipher the nanoscale arrangement of such biological complexes, considerable progress has been made in the past decades both in fluorescence-based super-resolution imaging (SRI) and transmission electron microscopy (TEM) ^6^. However, to meet the constantly improving spatial resolution offered by such advanced microscopy techniques, there is a need for small probes that label proteins of interest with high specificity, minimal distance to the target, strong penetration in restricted cellular compartments, and compatible with different microscopy modalities^7,8^.

As an alternative to traditional primary antibodies which are divalent, relatively large, and can be difficult to generate with good selectivity to target proteins, several single-chain antibodies or recombinantly expressed nanobodies have recently been engineered against either endogenous cytoskeletal or synaptic proteins ^9–11^, primary antibody IgGs ^11^, or popular tags like GFP, HA, or ALFA ^12–14^. These nanobodies can be chemically conjugated to a variety of fluorophores, allowing for multicolor detection in SRI techniques such as STORM or PAINT ^4,15–17^. Other protein probes including tetrameric or monomeric streptavidin bound to enzymatically biotinylated proteins, can be used as orthogonal labels in combination with nanobodies ^4^. Finally, recombinant proteins can be fused to enzymes such as SNAP or Halo tags followed by covalent coupling to their respective fluorescent ligands.

Whereas fluorescence-based SRI generates localization maps of specific proteins with little reference to sub-cellular structures, EM combined with heavy metal staining at room temperature, or performed at cryogenic temperature (CryoEM), provides exquisite identification of cell membranes and organelles ^18^, but is usually inferior in labeling individual proteins. Conventional immunogold techniques relying on antibodies conjugated to 5 or 10 nm gold particles ^19–21^, give mosaic, incomplete labeling of native proteins. Alternatively, recombinant proteins can be fused to peroxidases such as HRP, APEX, VIPER, or miniSOG and imaged after 3,3’-diaminobenzidine (DAB) reaction ^22–24^, but this yields a fairly uniform labeling where discrete protein clusters cannot readily be identified. Further developments for correlative light-electron microscopy (CLEM) have consisted in fusing APEX2 to a GFP nanobody to label GFP-tagged proteins ^25^, or in designing antibody or streptavidin fluoronanogold conjugates where the exact probe-to-protein stoichiometry is ill-known ^26,27^.

Thus, there is room for improved protein labeling strategies towards dual light and electron microscopy, which would allow for the detection of individual proteins by EM in relation to the nanodomains identified in SRI, and to sub-cellular morphology such as pre- and post-synaptic compartments. In light of current tools limitations, our goal was to design robust probes of small size, monovalency and strict 1:1 coupling to gold nanoparticle and fluorescent dye. We benchmarked our strategy on the GFP-Binding Protein (GBP), a widely used a nanobody that binds GFP and related derivatives with nanomolar affinity ^14,28^; GBP is a small monomer (∼3 nm, ∼10 kDa), easily produced in bacteria and broadly applied in SRI for (G)FP-tag detection ^4,15^. Strict stoichiometric coupling to a single monofunctional 1.4 nm-gold nanoparticle was achieved by keeping a unique - primary-amino group for robust activated ester chemistry. Further access to fluorescence imaging modalities was achieved by incorporation of an ALFA epitope tag, allowing its subsequent recognition with the secondary corresponding single domain antibody conjugated to an organic fluorophore. We applied this strategy to the visualization and counting of synaptic proteins in heterologous cells and neurons. This new probe specifically recognizes (G)FP-tagged proteins with a 1:1 stoichiometry, penetrates well into cell-cell contacts including synapses, and is compatible with both SRI and TEM. It should be applicable to study the localization of a wide range of (G)FP-tagged proteins in neuroscience and beyond.

## Results

### Engineering a nanobody probe with stoichiometric nanogold and fluorescent dye functionalization

We favored a strategy that relies on both natural amino acids and commercially available reagents to facilitate its implementation in as many laboratories as possible **(Fig. 1A)**. We first modified amino groups on GBP to ensure targeted conjugation of the gold nanoparticle. To enforce strict 1:1 coupling, we engineered the GBP to present a single amino group on a unique lysine, enabling direct amide formation to a monofunctional 1.4 nm N-hydroxysuccinimide (NHS)-gold nanoparticle. Based on the GFP/GBP complex (PDB 3K1K) ^29^, which shows no lysines in the GBP paratope **(Supp Fig. 1)**, we replaced all native lysines with arginines to preserve the surface charge distribution without impairing binding. In parallel, we installed a single lysine at the GBP C-terminus to provide a unique amino group, as the V_H_H N-terminus lies near the paratope and is unsuitable for attaching bulky moieties. To avoid the creation of an N-terminal amino group, we preserved the initiator formyl-methionine by placing an aspartate immediately after the start methionine, thereby preventing methionine aminopeptidase processing ^30^. We refer to the engineered GBP as GBP* **(Fig. 1B)**.

**Figure 1.**
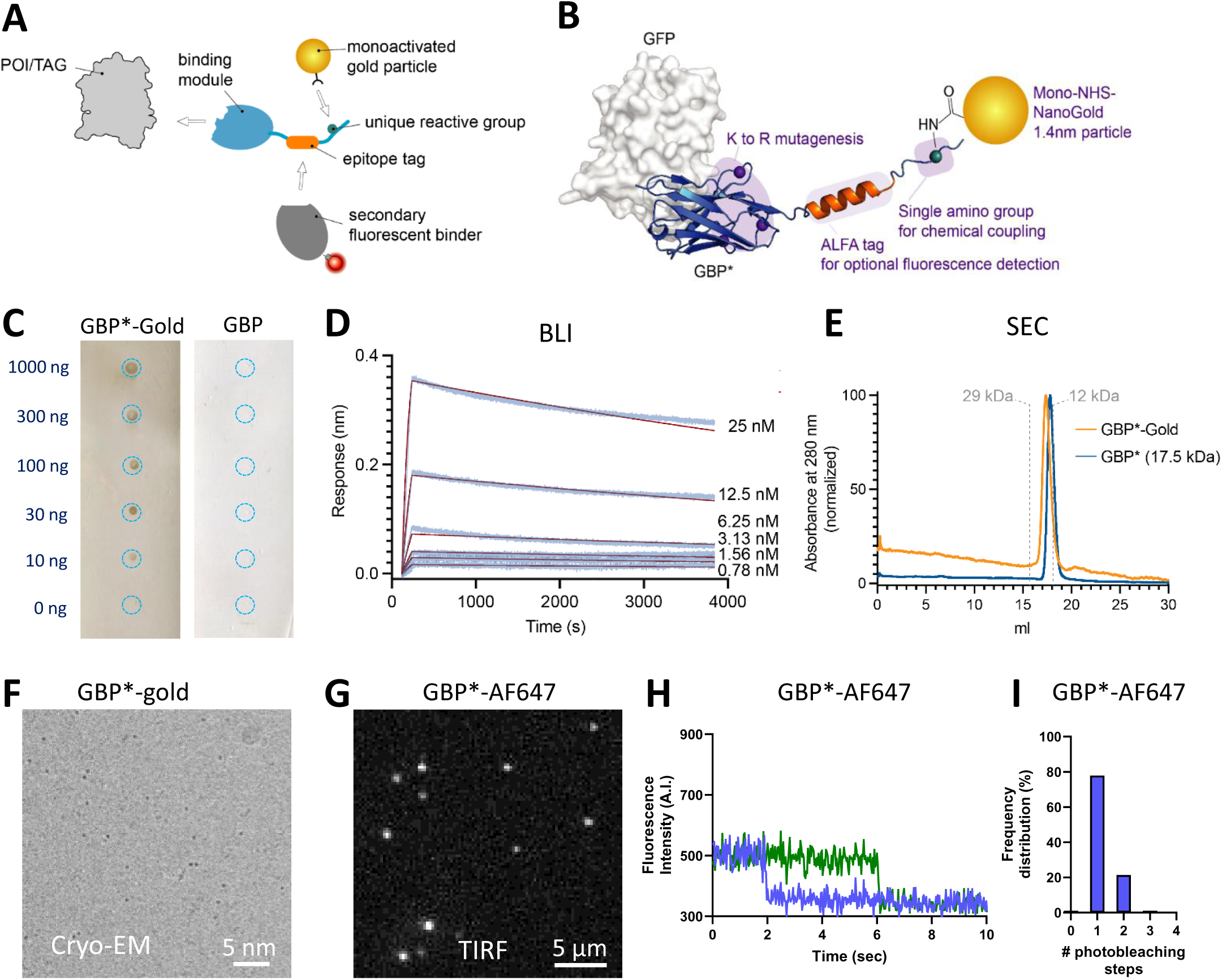
Engineering and in vitro characterization of the GBP*-gold. **(A)** Design of multimodal CLEM probes. **(B)** Implementation of the strategy to the GFP binding protein (GBP). The GBP* molecular model was generated by Alphafold3 and aligned to the GBP/GFP structure (PDB: 3K1K). **(C)** Dot blots with different amounts of GFP, labelled with GBP*-gold or unconjugated GBP, respectively, and silver enhanced. **(D)** Binding of purified GBP* to the EGFP target with BLI. Each line represents dilutions of the EGFP target (by 1:2). The red line show curve fits of raw data using the 1:1 global fitting model (K_D_ = 7.5 10^-10^ M, k_on_ 9.4 10^4^ 1/Ms, k_off_ = 7.1 10^-5^ 1/s). **(E)** Size-exclusion chromatography (SEC) profiles of the purified GBP* before and after coupling to the 1.4 nm gold particle, demonstrating single peaks consistent with the 1:1 conjugation. **(F)** Cryo-EM image of GBP*-gold on an EM grid, showing monodispersity of the conjugates. **(G, H)** Representative images of sparse GBP*-AF647 or GBP-AF647 molecules immobilized on a coverslip and imaged under TIRF illumination. **(I)** Fluorescence intensity over time of two randomly selected GBP*-AF647 molecules. Most molecules photobleach in one step, consistent with a 1:1 coupling of AF647 to the GBP*. Occasionally, two photobleaching steps are observed. **(J)** Histogram of photobleaching steps computed for 276 individual molecules over 14 acquisition sequences.

To establish a second, orthogonal route for installing SRI-compatible fluorescent dyes, we chose an indirect strategy. Although cysteines are often used orthogonally to lysines, this approach is limited by the lack of commercial alternatives to maleimide reagents, the instability of the maleimide–cysteine linkage^31,32^, and the need for extra post-conjugation purification. Moreover, directly coupling both a gold particle and an organic dye to a small domain such as a V_H_H can compromise solubility and increase nonspecific binding. Instead, we implemented a tag/binder system to confer fluorescence while minimizing the burden on the probe and enabling post-hoc modality changes. Accordingly, we appended a 12 aa long ALFA-tag at the GBP* C-terminus, leveraging its short α-helical, fixation-resistant sequence, absence of lysines, and subnanomolar affinity^13^.

### The GBP* probe is monofunctional, monodisperse, and retains binding to GFP in vitro

The resulting construct was produced in bacteria and functionality of the GBP* variant was first investigated with either non-conjugated, organic dye- or gold-conjugated nanobodies. We first confirmed that the modified nanobody retained its binding affinity to purified GFP in vitro, as determined qualitatively by dot blot **(Fig. 1C)**, and quantitatively by BLI with a 0.75 nM affinity very similar to reported affinities for the non-modified nanobody ^29^ **(Fig. 1D)**. Analysis of GBP* and the GBP*-gold conjugate by size-exclusion chromatography revealed in both cases monomeric and homogeneous proteins with a shorter retention time for the gold conjugated sample consistent with a single coupling event **(Fig. 1E)**. When the purified GBP*-gold preparation was spread on an EM grid, we observed fairly monodisperse nanogold particles, indicating no sign of aggregation **(Fig. 1F)**. To check that the engineered antibody was indeed monoreactive, it was conjugated to an NHS-ester Alexa Fluor 647 dye (in place of the gold nanoparticle), and individual photobleaching steps were quantified upon laser illumination of a diluted suspension of GBP*-AF647 immobilized on a coverslip **(Fig. 1G,H)**. We observed a vast majority (80%) of single photobleaching steps **(Fig. 1I,J)**, confirming the fact that the GBP* is essentially conjugated to a single chemical moiety.

### The GBP*-gold conjugate specifically binds GFP-tagged proteins in heterologous cells

We then assessed the ability of the GBP*-gold conjugate to recognize GFP-tagged proteins in cell membranes. We electroporated COS-7 cells with the synaptic adhesion molecule neurexin 1β (Nrxn1β) bearing an extracellular GFP-tag at the N-terminus ^33^ **(Fig. 2A)**. After 36 hrs to allow for protein expression and targeting to the cell surface **(Fig. 2B)**, cells were sequentially incubated live with GBP*-gold, then with Nb-ALFA conjugated to AF647 **(Fig. 2A)**. Cells expressing GFP-Nrxn1β were specifically labelled with Nb-ALFA-AF647, demonstrating that the GBP*-gold was bound to surface GFP-Nrxn1β molecules, and recognized by Nb-ALFA-AF647 **(Fig. 2C)**. After recording their position on the microscope, cells were fixed and nanogold particles were silver enhanced for 40 min. A corresponding signal was detected in differential interference contrast (DIC) on cells expressing GFP-Nrxn1β positive cells, but not on untransfected neighboring cells **(Fig. 2D**, asterisk). Overall, there was a positive correlation between the silver enhanced GBP*-gold signal in DIC and the GFP fluorescence signal which varies from cell to cell according to the GFP-Nrxn1β expression level **(Fig. 2E)**. Finally, COS-7 cells electroporated with GFP-Nrxn1β were live labeled with GBP*-gold, and processed for TEM after fixation and mild silver enhancement (10 min). We detected a pearl-like arrangement of single nanoparticles along the cell contours, which was absent from non-electroporated cells **(Fig. 2F,G)**, thus revealing the specific detection of GFP-Nrxn1β expressed at the cell membrane. For these applications, COS-7 cells were plated on etched coverslips bearing an internal reference coordinate system allowing us to track the Nrxn1β positive cells in fluorescence microscopy, then find them later in TEM after resin embedding and ultrathin sectioning **(Fig. S3)**. As a powerful additional feature of this labelling strategy, the use of the Nb-ALFA-AF647 gave us the opportunity to perform dSTORM experiments on GFP-Nrxn1β expressing COS-7 cells, using the AF647 fluorophore as a stochastic emitter **(Fig. S2A)**.

**Figure 2.**
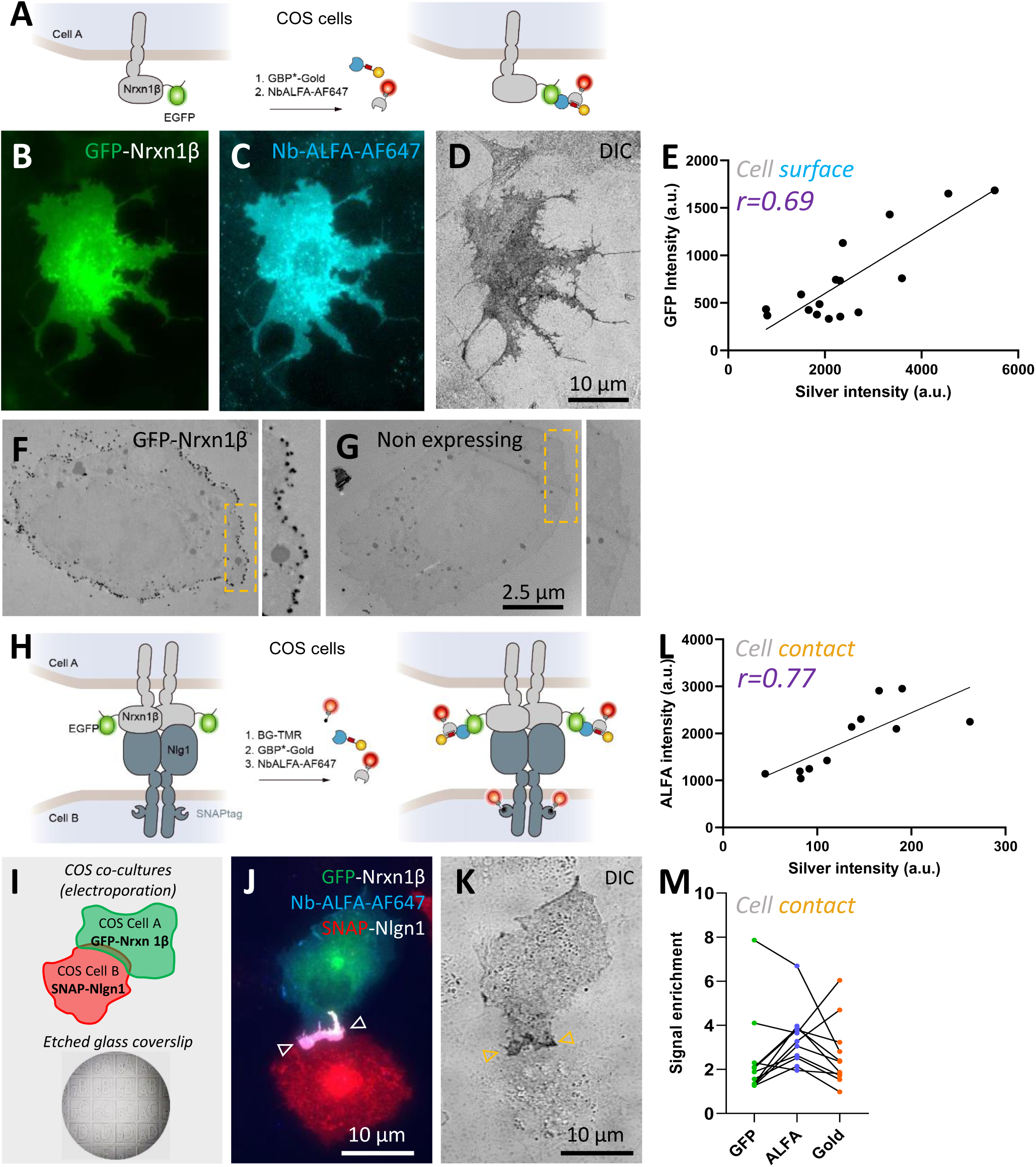
The GBP*-gold specifically labels cells expressing GFP-tagged proteins and penetrates in cell-cell contacts. **(A)** Schematics of GFP-Nrxn1β expression and labelling in COS-7 cells. **(B)** Representative epifluorescence image of a COS-7 cell expressing GFP-Nrxn1β plated on an etched coverslip. **(C)** Cells were sequentially labeled with GBP*-gold and Nb-ALFA-AF647. **(D)** Cells were fixed, silver enhanced, and the same cell was imaged in DIC. Note that the GFP-Nrxn1β negative cell in the upper right corner does not show signals of Nb-ALFA-AF647 or silver. **(E)** Correlation between the GFP-Nrxn1β signal and the silver precipitate, dots representing individual cells. **(F, G)** TEM of COS-7 cells with or without GFP-Nrxn1β expression, respectively, labeled with GBP*-gold and silver enhanced. **(H, I)** Diagrams of the orthogonal labelling of GFP-Nrxn1β and Nlgn1-SNAP in COS-7 cell contacts. **(J)** Image of a contact between two COS-7 cells, one expressing GFP-Nrxn1β (green) and the other expressing Nlgn1-SNAP (red) labeled with TMR-conjugated SNAP-cell substrate (red). Note the accumulation of the two adhesion proteins in the cell-cell contact (arrows). Cells were sequentially live labeled with GBP*-gold and Nb-ALFA-AF647 (cyan), then fixed and silver enhanced **(K)**. Note the accumulation of both the Nb-ALFA-AF647 and the silver precipitate in the adhesive contact, revealing penetration of the GBP*-gold, and no staining of the cell expressing Nlgn1-SNAP. **(L)** Correlation between Nb-ALFA-AF647 fluorescence intensity and silver-enhanced signal in the cell contact. Dots represent individual cell contacts. (**M**) Enrichment of GFP-Nrxn1β, Nb-ALFA-AF647, and silver precipitate in adhesive contacts, defined as the ratio between the level in the cell-cell contact, and the relative level in regions of the cells outside the contact.

### The GBP*-gold conjugate penetrates well in cell-cell contacts

To assess whether the GBP*-gold conjugate can penetrate tight cell-cell contacts, we mixed COS-7 cells expressing GFP-Nrxn1β together with cells expressing Nlgn1 fused to an intracellular SNAP tag **(Fig. 2H)**. This approach results in the formation of large lamellipodial cell-cell contacts that look like giant synapses, where Nrxn1β and Nlgn1 bind almost irreversibly to each other and accumulate over time ^33^ **(Fig. 2I)**. After staining the SNAP tag with the cell-permeant SNAP tag substrate conjugated to tetramethyl rhodamine (SNAP-Cell-TMR), we sequentially incubated the co-cultures with GBP*-gold, then with Nb-ALFA-AF647 **(Fig. 2J)**. We took fluorescence images, recorded the positions, then fixed the cells and silver enhanced the gold nanoparticles before taking DIC images **(Fig. 2K)**. The dark silver precipitate deposited on cells expressing GFP-Nrxn1β was stronger at contacts with cells expressing Nlgn1-SNAP and correlated well with the Nb-ALFA-AF647 signal **(Fig. 2L)**. The enrichment factor, taken as the ratio of the signal at the cell-cell contact divided by the respective signal on the cell surface outside the contact, was around 3, similar to the enrichment calculated from the GFP-Nrxn1β or the Nb-ALFA-AF647 signals **(Fig. 2M)**. This result demonstrates that the GBP*-gold penetrates readily into cell-cell junctions, without steric hindrance from the 1.4-nm nanogold particle, encouraging us to look at more complex sub-cellular structures such as neuronal synapses.

### The GBP*-gold conjugate detects endogenous GFP-βNrxn molecules in synapses

To assess whether the GBP*-gold conjugate could stain GFP-tagged molecules expressed at endogenous levels in synapses, we first used primary hippocampal cultures from a previously described knock-in (KI) mouse strain in which native Nrxn1β and Nrxn3β are tagged with GFP ^34^. At days in vitro (DIV) 18, we stained the neurons live with GBP*-gold, and processed the samples for TEM **(Fig. 3A)**. We identified silver-enhanced nanoparticles in the synaptic cleft of KI neurons **(Figs. & S4A)**, indicating that our strategy is sensitive enough to locate endogenous βNrxn molecules that are expressed only at very low copy numbers per synapse ^35–37^. As a negative control, there was no signal in cultures from wild-type mice treated similarly with GBP*-gold, attesting to the specificity of our approach **(Fig. 3C)**. In addition to the synaptic cleft association, we found endogenous GFP-βNrxns over pre-synaptic terminals and along the axons outside of synaptic protrusions **(Fig. S4B-C),** consistent with earlier results using a divalent anti-GFP antibody^34^. We quantified GFP-βNrxn molecules in a limited number of TEM images from our GBP*-gold labeled hippocampal KI cultures and observed that they are 2-3 fold more abundant on axons compared to the pre-synapses or the synaptic cleft **(Fig. 3D)**.

**Figure 3.**
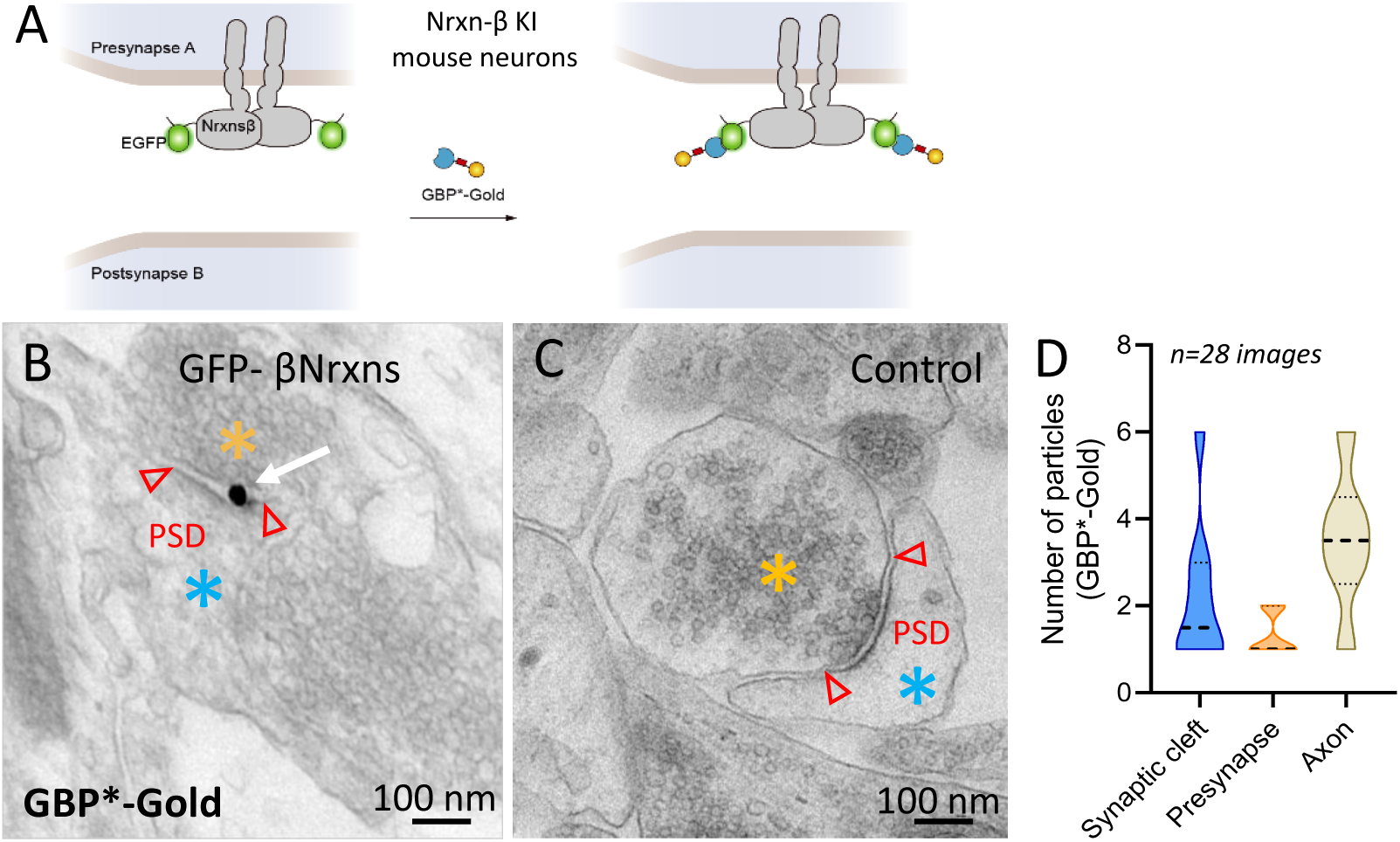
Visualization of endogenous β-Nrxns using the GBP*-gold. **(A)** Schematics of GFP-tagged, endogenous βNrxn labelling in neurons. **(B)** Representative TEM image of a synapse in neuronal cultures derived from KI mice (GFP-βNrxn KI) expressing GFP-Nrxn1β and GFP-Nrxn3β. Cultured neurons were live-labeled with GBP*-gold, then fixed and silver-enhanced before processing for TEM. The white arrow indicates βNrxn in a synaptic cleft between a presynaptic terminal (orange asterisks) filled with synaptic vesicles (SVs) and a postsynaptic compartment (blue asterisks). **(C)** Image showing the absence of labelling in control neurons from WT mice. (**C**) Quantitative analysis of the subcellular distribution of GBP*-gold particles in the synaptic cleft, pre-synapse, or axonal compartment. Data represent mean ± SEM and were compared by a Kruskal-Wallis test followed by Dunn’s multiple comparison tests (****P < 0.0001).

### The GBP*-gold conjugate stains overexpressed GFP-tagged AMPA receptors in synapses

To target an abundant and widely studied post-synaptic protein, we examined the localization of AMPA receptors with our new probe. Dissociated rat hippocampal neurons were initially electroporated with the AMPA receptor subunit GluA1 N-terminally tagged with SEP (SEP-GluA1), a recombinant protein that accumulates well in post-synapses ^38,39^ **(Fig. 4A)**. At DIV14, neurons plated on etched glass coverslips were incubated with GBP*-gold, then with Nb-ALFA-AF647 **(Fig. 4B,C)**. The two signals colocalized well **(Fig. 4E)**, indicating the penetration of the GBP*-gold in synapses. Neurons were then fixed, silver enhanced, and examined by light microscopy. A dark punctate staining was seen along the dendrites of SEP-GluA1 expressing neurons, with enrichment in dendritic spines **(Fig. 4D,E)**. Interestingly, the Nb-ALFA-AF647 allowed us to perform dSTORM observation of SEP-GluA1 prior to TEM **(Fig. S2B)**, revealing its nanoscale organization in synapses, where a few AMPA receptor nanodomains could be identified in each post-synapse **(Fig. 4F-H)**, as already reported ^2,3,5,40^. By normalizing the number of Nb-ALFA-AF647 localizations in synapses, to the number of localizations given by single nanobodies bound to the coverslip, we estimated a copy number of 23.7 ± 2.3 over-expressed GluA1-containing AMPARs in individual synapses (n = 43). We then processed the same samples for TEM **(Fig. S3)**, and observed a striking accumulation of silver-enhanced nanogold particles in the synaptic cleft **(Fig. 4I)**. The number of particles (13.3 ± 2.4) was lower than the numbers found by dSTORM (23.7 ± 2.3) (**Fig. 4J**), likely due to the fact that ultrathin sectioning underestimates the actual number of receptors in a whole PSD.

**Figure 4.**
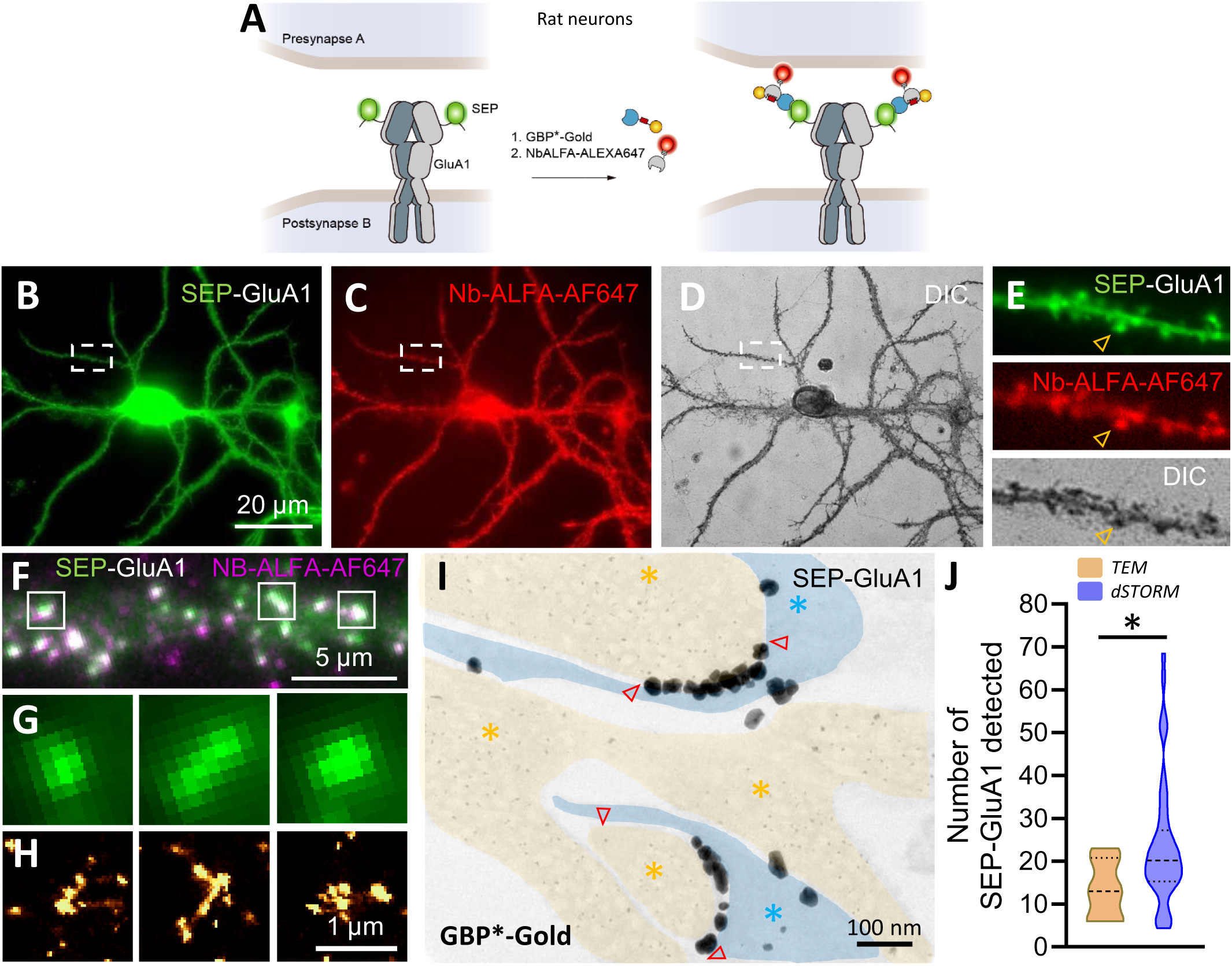
GBP*-gold staining of recombinant GFP-tagged neuronal proteins in synapses. **(A)** Diagram of the labelling of recombinant SEP-GluA1 expressed in rat hippocampal neurons. **(B-D)** Representative images of a neuron expressing SEP-GluA1 (green) sequentially labeled with GBP*-gold and Nb-ALFA-AF647 (red), then fixed and silver enhanced (dark). **(E)** Zoom on a dendritic segment, with the same color code. **(F)** Two-color epifluorescence image of a dendritic segment from a neuron expressing SEP-GluA1 (green) and labelled with Nb-ALFA-AF647 (magenta). **(G)** Zoom on three square synaptic regions selected from (F), showing the punctate SEP-GluA1 signal (green). **(H)** Corresponding localization maps of Nb-ALFA-AF647 performed in dSTORM, showing AMPA receptor nanodomains. **(I)** Representative TEM image of a synapse showing the accumulation of silver-enhanced GBP*-gold in the synaptic cleft, delimited by arrows. Pre- and post-synaptic compartments are shown in yellow and blue, respectively. **(J)** Number of synaptic SEP-GluA1 molecules measured either from the Nb-ALFA-AF647 localizations in dSTORM images, or from the GBP*-gold nanoparticles in TEM.

### The GBP*-gold conjugate labels endogenous GFP-tagged AMPA receptors in synapses

Finally, to stain AMPA receptors expressed at endogenous levels, we used genome editing based on a CRISPR/Cas9 strategy, called TKIT, consisting in replacing endogenous GluA2 subunits by SEP-tagged counterparts in post-mitotic cells such as neurons, using specific guide RNAs ^41^ **(Fig. 5A)**. In practice, we dissociated hippocampal cells from transgenic mouse pups ubiquitously expressing Cas9 ^42^, cultured them in spheroids, and infected the spheroids with AAVs packaging gRNAs for the knock-in of SEP-GluA2 in single neurons, before processing the samples at DIV14. Such neurospheres provide a more physiologically relevant, tissue-like environment than primary cultures made on glass coverslips and are easily processed for either confocal microscopy or TEM ^43^. Several SEP-GluA2-positive neurons were identified per spheroid by confocal microscopy, each with plenty of dendritic spines enriched in SEP-GluA2 signal **(Fig. 5B,C)**. After live labeling with GBP*-gold, fixation, and silver enhancement, spheroids were recovered by sedimentation prior to resin embedding, and 3D samples allowed the collection of many ultrathin sections for the observation of synapses in TEM **(Fig. 5D)**. Endogenously expressed SEP-GluA2 containing AMPA receptors were labelled with many gold nanoparticles in the synaptic cleft, often forming discrete sub-domains **(Fig. 5E)**. There were no nanoparticles in synapses when spheroids were infected with a control AAV IRES-GFP, revealing again the staining specificity of the GBP*-gold in this sample.

**Figure 5.**
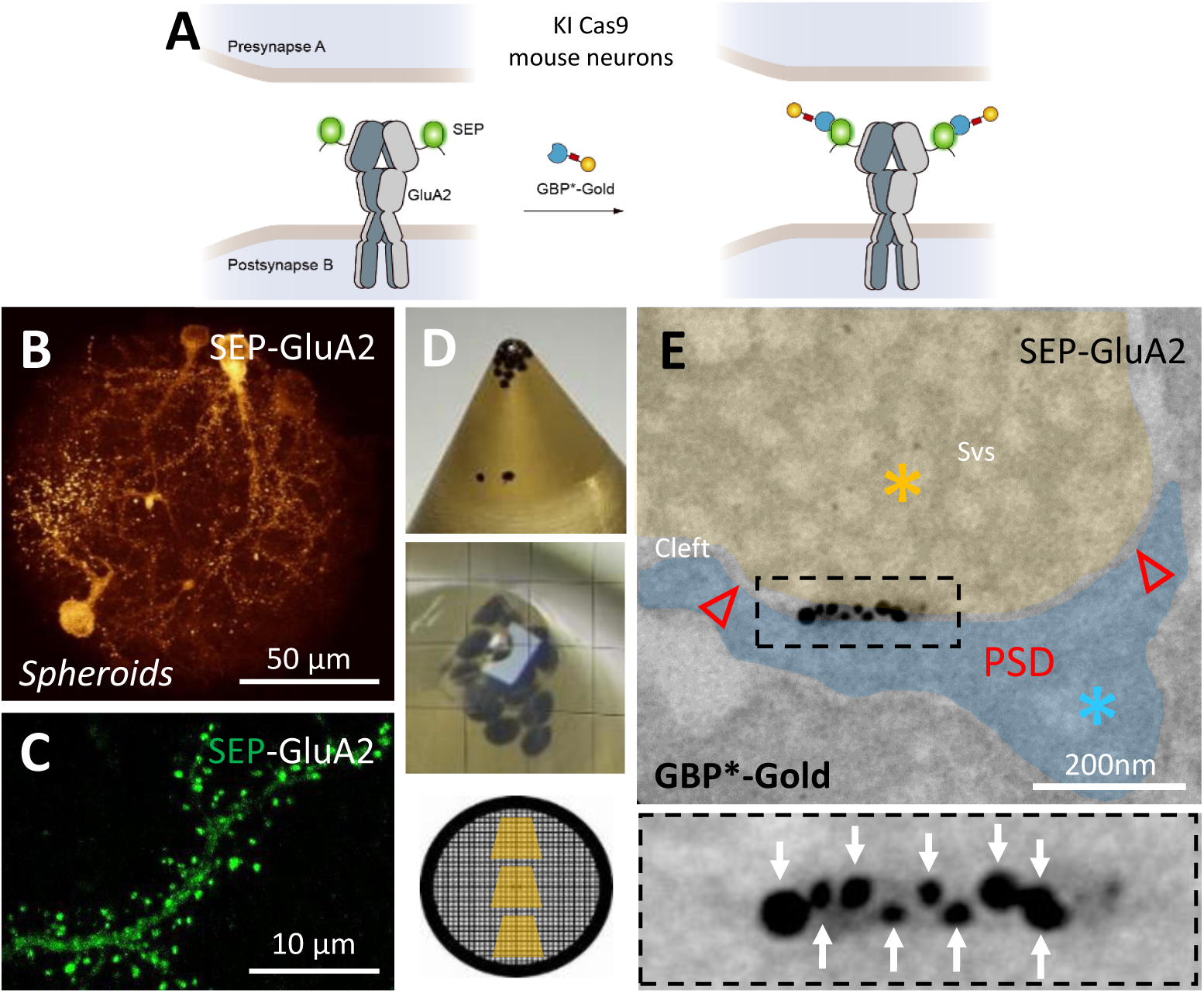
GBP*-gold labelling of genome-edited SEP-GluA2 in neurons. **(A)** Schematics of the labelling of endogenous SEP-GluA2 in neurons from Cas9 mice. **(B)** Maximal 2D projection confocal image of an entire neuronal spheroid generated from hippocampal cells of Cas9-expressing mice, following genome editing of endogenous SEP-GluA2 (gold). **(C)** Zoom on a dendritic segment showing enrichment of SEP-GluA2 (green) in dendritic spines. **(D)** Sample preparation workflow for EM. (Top) representative image of neuronal spheroids sedimented at the bottom of a tube and embedded in resin; (Middle) Block trimming; (Bottom) Ultrathin sections (70 nm) subsequently cut and collected onto EM grids. **(E)** TEM image of a synapse showing the accumulation of silver-enhanced GBP*-gold particles in the synaptic cleft, that specifically label native SEP-GluA2 receptors. The inset zooms on an AMPA receptor nanodomain with high nanoparticle density.

## Discussion

To probe the internal organization of cellular junctions such as neuronal synapses by super-resolution microscopy, there is a need for small, multimodal probes able to access specific protein targets in narrow contact zones filled with a dense multi-molecular network. In this context, we developed here a bifunctional probe that labels GFP-tagged proteins with high selectivity for both light and electron microscopy. The key design principle was stoichiometry control: by re-engineering the GFP nanobody to present a single primary amine, we achieved NHS-mediated attachment of exactly one 1.4-nm gold particle per binder. This resolves the compositional uncertainty inherent to conventional fluoronanogold reagents, where multiple lysines on carriers such as Fab fragments or streptavidin permit variable dye-to-gold ratios. Because the 1.4-nm gold is monofunctional, the resulting complex enforces one gold per target and one epitope engagement, allowing protein copy-number estimation from EM particle counts. Orthogonally, a C-terminal ALFA tag provides flexible SRI compatibility: using a second anti-ALFA nanobody, one can introduce dSTORM-suitable dyes (AF532/AF647) ^4,44,45^, DNA handles for DNA-PAINT^16^, or STED-optimized fluorophores (AF594, ATTO647N) ^46^ without perturbing the gold conjugate.

The GFP nanobody is around 3 nm in size and is expected not to be affected by the coupling to a 1.4 nm gold nanoparticle. Indeed, the GBP*-gold conjugate retained a strong affinity to purified GFP in vitro and to GFP-tagged proteins expressed on the cell surface. Moreover, this probe penetrated readily into tight cell-cell contacts and in the synaptic cleft that are around 30 nm accross ^47^. This is a strong advantage compared to other protein tracking approaches that have relied on the use of larger probes such as gold nanoparticles or Quantum dots coated with proteins, where the overall size and multivalence of those conjugates can lead to steric hindrance and make them hard to access narrow cell-cell or cell-matrix contacts ^48,49^. Based on our own experience, we found that there is a sharp size cut-off around 5 nm above which probes experience difficulty accessing such contacts, as demonstrated by comparing the penetrability of monomeric streptavidin (3 nm) versus anti-biotin antibody (12 nm) ^4,50^.

Leveraging on the 1:1 stoichiometry between target and probe, we could estimate the copy numbers of individual synaptic proteins including Nrxns and AMPA receptor subunits in the synaptic cleft. We found 1 to 2 nanoparticles bound to native βNrxn molecules in the pre-synapse. Since ultrathin sections are 70 nm thick, this number should be multiplied by a factor of around 3 to match the width of the synaptic cleft. Such an estimate of 3-6 βNrxn molecules is compatible with dSTORM staining of endogenous Nrxns performed using an HA-tagged Nrxn1 knock-in mouse strain ^37^. Those numbers appear quite low given the large repertoire of known Nrxn binding partners in synapses ^51^, but may be compensated by the expression of other Nrxn isoforms ^35,52^. In contrast, and as expected from the literature, we found larger numbers of post-synaptic GFP-tagged AMPA receptor subunits, i.e. around 24 copies of overexpressed GFP-GluA1 subunits by dSTORM, and 13 copies by TEM (i.e. to be multiplied by ∼ 3 due to PSD sectioning). Native SEP-GluA2 subunits were visually as abundant as over-expressed GFP-GluA1, potentially because they assemble into both populations of GluA1/GluA2 vs GluA2/GluA3 heterotetramers in hippocampal neurons ^53,54^. Furthermore SEP-GluA2 subunits formed nanodomains, as already identified with antibodies against GluA2 in rat primary neurons ^2^, or with streptavidin-nanogold conjugates on hippocampal cultures from a knock-in mouse where endogenous GluA2 carries a biotinylatable tag ^26^, although in the latter experiments there is always a risk of protein cross-linking because of probe multivalence. The virtually background-free labeling method developed here, along with the supposedly 1:1 stoichiometry of label:molecule in the GBP*-gold, will allow reliable estimations, for example, of actual copy numbers of synaptic proteins in the future that will go beyond conventional immunoEM with antibodies.

As perspectives, because silver enhancement is based on a nucleation process which is hard to control and can lead to an overall dispersity of aggregate size, one might want to image bare 1.4 nm gold particles bound to nanobodies, e.g. using cryoEM and ideally in 3D tomography, as was done recently with Fabs to AMPA receptors conjugated to 2-nm gold particles ^55^. Moreover, our monofunctional gold conjugation strategy could be applied to probes including other nanobodies or monomeric streptavidin ^4^ for multicolour EM using gold probes of different sizes (e.g. 1.4 and 3 nm). By combining this approach with the crossing of GFP-tagged and bAP-tagged knock-in mice ^26,56^, one might aim at simultaneously visualizing streptavidin-bound biotinylated proteins such as Nlgn1 or GluA2 subunits, in proximity to nanobody-bound GFP-tagged proteins such as Nrxns in the synaptic cleft at an unprecedented single-molecule resolution level, with the aim to shed more light on the synaptic nanocolumn paradigm ^1^.

## Materials and Methods

### Animals

Pregnant Sprague-Dawley rat females were purchased from Janvier Labs (Saint-Berthevin, France). The H11-Cas9 mouse strain B6J.129(Cg)-*Igs2^tm^*^1*.1(CAG-cas9*)Mmw*^/J was imported from Jackson Labs (ref : 028239) and raised in the animal facility at the University of Bordeaux. To avoid genetic drift, mice were back-crossed with wild-type (WT) mice every 10 generations. Triple βNrxn knock-in mice (βNrxn KI) are available from Jackson Labs (B6;129-Nrxn3tm2Sud Nrxn1tm2Sud Nrxn2tm2Sud/J) and were previously described ^34,57^. These floxed KI mice carry epitope tags after the βNrxn-specific signal peptide sequence, resulting in expression of EGFP fused N-terminally to the mature sequence of Nrxn1β and Nrxn3β (GFP-Nrxn1β and GFP-Nrxn3β), and of an HA tag to Nrxn2β. All animals were handled and killed according to European ethical rules, with the agreement of the local ethics committees from the University of Bordeaux (EC50) and University of Münster, and approved by the Landesamt für Natur, Umwelt und Verbraucherschutz (LANUV, NRW, Germany), license numbers 84-02.05.20.11.209 and 84-02.04.2015.A423.

### Constructs

GFP-Nrxn1β and SEP-GluA1 were previously described ^35,38,39^. HA-Nlgn1 ^58^ was a kind gift from P. Scheiffele (Biozentrum, Basel, Switzerland). HA-Nlgn1-SNAP was made by introducing the SNAP tag in place of the GFP tag 22 aa upstream of the C-terminus in the previously described plasmid HA-Nlgn1-GFP ^59^. The engineered GBP (GBP*, K to R mutation and installation of a single C-terminal lysine) was obtained by gene synthesis (Eurofins) with codons optimized for E. coli expression. The synthetic gene was subcloned via NdeI and XhoI restriction sites into a home-made vector, pIGc, with a C-teminal decahistidine tag. The ALFA tag was added between the nanobody C-terminus and the unique lysine group by IVA cloning ^60^. The ALFA nanobody was obtained by gene synthesis (Eurofins) as a variant with mutation of the surface nanobody lysine groups into arginines. The synthetic gene was subcloned via BamHI and HindIII restriction sites into a home-made vector with a N-terminal 14-Histidine tag, in frame with a maltose-binding protein followed by a NEDD8 (*Brachypodium distachyon*) domain, and a C-teminal additional lysine. EGFP was subcloned into our pIGb vector ^61^ using NdeI and XhoI restriction sites to be produced in bacteria with a C-terminal biotin acceptor peptide site and decahistidine tag. The bdNEDP1 protease (pDG02583) was a gift from D. Görlich (Addgene plasmid # 104129) ^62^.

### Engineering and production of GFP and ALFA nanobodies

The modified GBP*, GBP, and ALFA-tag nanobodies were produced as described previously ^4^. Briefly, constructs were expressed in E. coli BL21(DE3) by auto-induction at 16°C and purified under native conditions by Ni²⁺-affinity *via* their polyhistidine tags. The ALFA-tag nanobody was further processed with in-house produced bdNEDP1 protease ^63^. Proteins were then purified by size-exclusion chromatography in PBS and concentrated to ∼1 mg·mL⁻¹. Functionalization was achieved by coupling either 3–6 equivalents of Alexa Fluor 647 (Thermo Fisher) or 1.1 equivalents of 1.4-nm monofunctional gold nanoparticles (Mono-Sulfo-NHS-Nanogold®, Nanoprobes), both in NHS-ester form. Excess free dye was removed on a desalting column, and probe-conjugated nanobodies were further purified by size-exclusion chromatography. Conjugates were concentrated; AF647-nanobodies were flash-frozen and stored at −80 °C, whereas Nanogold-conjugated GBP* was stored at 4 °C for several weeks. Biotinylated EGFP was produced in home-made E. coli BL21(DE3) cells containing the pACYC-mCh-BirA plasmid as previously described ^61^. For analytical size-exclusion chromatography, representative GBP*-gold conjugates and GBP* were injected into a Superdex 200 Increase 10/300 GL column on an AKTA purifier LC system. Samples were run in PBS at 0.75 ml/min and monitored at 280 nm.

### Biolayer interferometry

Bio-layer interferometry (BLI) measurements were acquired on an Octet R8 (Sartorius) and analyzed with the instrument software. To assess the affinity of the engineered GBP*, C-terminally biotinylated EGFP (∼50 nM in PBS, 0.05% Tween-20, 0.1% biotin-free BSA) was immobilized on streptavidin (SA, Sartorius) biosensors to ∼0.5 nm response. GBP analyte was prepared by diluting the concentrated stock into the same buffer. After establishing a baseline, association was recorded by dipping sensors into GBP at 25, 12.5, 6.25, 3.13, 1.56, or 0.78 nM, followed by dissociation in buffer. Global kinetic fitting with a 1:1 model was used to determine the K_d_ across the dilution series.

### COS-7 cell culture and electroporation

COS-7 cells from the ECACC purchased via Sigma-Aldrich, Acc Nc 87021302) were cultured in DMEM (Eurobio) supplemented with 1% glutamax (Sigma-Aldrich, #3550-038), 1% sodium pyruvate (Sigma-Aldrich, #11360-070), 10% Fetal Bovine Serum (Eurobio). Cells were thawed from frozen vials at passage +4, and maintained up to passage 20. Cells were regularly tested negative for mycoplasma, using the MycoAlert detection kit (Lonza, #LT07-218). At confluence, cells from a T25 flask were treated for 5 min with Trypsin-EDTA at 37°C. Detached cells were harvested in supplemented DMEM and centrifuged at 1,000 rpm for 5 min. After discarding the supernatant, the cell pellet was resuspended in 200 µL of cell line V electroporation solution. Half of the cells were mixed with 5 µg GFP-NRrxn1β and the other half with 5 µg Nlgn1-SNAP. Samples were placed in two separate cuvettes and exposed to electroporation (Amaxa system, Lonza) using the COS-7 ATCC program. Electroporated cells were resuspended in 1 mL culture medium, and 50 µL of each suspension was added to each well of a 12-well plate containing 1 mL of culture medium per well and an 18 mm glass coverslip at the bottom. In some experiments, COS-7 cells were seeded on photoetched square coverslips of 18 x 18 mm (Electron Microscopy Science, # 72264-18) for an easy tracking of cells of interest. In that case, cells were cultured in a 6-well plate with 2 mL culture medium per well. Cells were left in a humidified 37°C - 5% CO_2_ incubator for 36-48 hrs before the experiment.

### Dissociated rat neuronal cultures and electroporation

Dissociated neurons were prepared from E18 rat embryos as previously described ^46,64^. Right after the dissociation, 300,000 cells were resuspended in 100 µL of P3 Primary Cell solution X kit (Lonza, # V4XP-3024), mixed with 5 µg SEP-GluA1 DNA, placed in a cuvette, and electroporated with the 4D-Nucleofactor (Lonza) using the High Viability program. Electroporated neurons were resuspended in Minimal Essential Medium (Thermo Fisher Scientific, #21090.022) supplemented with 10% Horse serum (Invitrogen) (MEM-HS), using the plastic pipette provided by the manufacturer, and plated on 18 mm glass coverslips previously coated with 1 mg.mL^-1^ polylysine (Sigma-Aldrich, #P2636) overnight at 37°C. To promote neuronal development, 75,000 non electroporated neurons were added to the dish. In some experiments, neurons were seeded on photoetched coverslips (Electron Microscopy Science, # 72264-18) similarly coated with polylysine. Three hours after plating, coverslips that contain each three paraffin dots acting as spacers, were flipped onto 60 mm dishes containing 15 DIV rat hippocampal glial cells cultured in Neurobasal plus medium (Gibco Thermo Fisher Scientific, #A3582901) supplemented with 2 mM glutamine and 1x B27^TM^ plus Neuronal supplement (Gibco Thermo Fisher Scientific, #A3582801). Astrocyte feeder layers were prepared from the same embryos, plated at 30,000 cells per 60 mm dish previously coated with 0.1 mg.mL^-1^ polylysine and cultured for 14 days in MEM containing 4.5 g.L^-1^ glucose, 2 mM L-glutamax (Sigma-Aldrich, #3550-038) and 10% horse serum. Ara C (Sigma-Aldrich, #C1768) was added after 3 DIV at a final concentration of 2 µM. Neurons were cultured for 14 days in a 37°C - 5% CO_2_ incubator.

### Cell labeling with nanobodies and observation by epifluorescence microscopy

COS-7 cell co-cultures dually expressing GFP-Nrxn1β and Nlgn1-SNAP were first labeled for 30 min with SNAP-Cell® TMR-Star (New England Biolabs, # S9105S, stock solution 0.6 mM in DMSO) diluted 1:200 in 1 mL culture medium, then rinsed twice with 1 mL warm culture medium. GFP-Nrxn1β was further live labelled for 10 min at 37°C with GBP*-gold diluted 1:100 in 200 µL Tyrode solution (15 mM D-glucose, 108 mM NaCl, 5 mM KCl, 2 mM MgCl_2_, 2 mM CaCl_2_ and 25 mM HEPES, pH = 7.4, 280 mOsm) supplemented with 1% Bovine serum Albumin (BSA, A3059-50G). Cells were rinsed twice with 1 mL warm Tyrode’s solution, then incubated live for 10 min at 37°C with the Nb-ALFA-AF647 diluted 1:100 in 200 µL Tyrode’s solution containing 1% BSA. Incubations with nanobodies were made by placing the coverslip upside down onto a piece of parafilm containing the 200 µL nanobody dilution. Similarly, DIV14 neurons expressing SEP-GluA1 were sequentially incubated with GBP*-gold, then Nb-ALFA-AF647. Stained cultures were then mounted in a dedicated Inox observation chamber (Ludin, LSI), and observed live under an inverted epifluorescence microscope (Nikon Eclipse TiE) surrounded by a thermostated Plexiglas box at 37°C, and equipped with a 60x/1.45 NA objective and filter sets for EGFP (Excitation: FF01-472/30; Dichroic: FF-495Di02; Emission: FF01-525/30); TMR-Star (Excitation: FF01-543/22; Dichroic: FF-562Di02; Emission: FF01-593/40); and AlexaFluor 647 (Excitation: FF02-628/40; Dichroic: FF-660Di02; Emission: FF01-692/40) (SemROCK). Images of 10-15 electroporated cells were acquired using an sCMOS camera (PRIME 95B, Photometrics) driven by the Metamorph® software (Molecular Devices), using exposure times of 100-500 ms adjusted to optimize the fluorescence signal in each channel. After recording the positions of the cells of interest, cells were fixed for 10 min in 4% paraformaldehyde (PFA) -4% sucrose in PBS at room temperature then kept in PBS at 4°C until dSTORM acquisitions, or exposed to silver enhancement for 40 min (LI Silver, Nanoprobes Inc) in the dark before being placed back onto the microscope. Differential interference contrast (DIC) at 50 ms exposure was then used to image the silver precipitate that had formed on cells labeled with GBP*-gold. The etched patterns on the coverslips were used to track the exact same cells that had been previously imaged in epifluorescence. The average fluorescence and DIC signal intensities were measured on the cells of interest using a thresholding procedure under Metamorph. For TEM, the coverslips containing COS-7 cells or neurons expressing GFP-tagged proteins and labeled with GBP*-gold were fixed overnight at 4 °C in PBS with 4% PFA-sucrose and 0.2% glutaraldehyde.

### dSTORM experiments and image reconstruction

The coverslips containing COS-7 cells or neurons expressing GFP-tagged proteins and labeled with Nb-ALFA-AF647 were mounted in a Ludin chamber in an oxygen-scavenging imaging buffer: Tris-HCl buffer (pH 7.5), containing 10% glycerol, 10% glucose, 0.5 mg.mL^-1^ glucose oxydase (Sigma-Aldrich, #G2133), 40 mg.mL^-1^ catalase (Sigma-Aldrich, #C100, 0,1% w/v) and 50 mM β-mercaptoethylamine (MEA, Sigma-Aldrich, #M6500) ^44^. Nano-diamonds (Adamas Nanotechnologies, #ND-NV140 nm) were added for 5 min to the samples for later registration of images and lateral drift correction. Chambers were then sealed using a second glass coverslip (24 mm diameter) and placed under the same microscope described above for live-labeling, which is further equipped with a perfect focus system preventing drift in the z-axis. A four-color laser bench (405/488/561 nm lines, 100 mW each, Roper Scientific; and 647 nm line, MPB Communications Inc.) is connected through an optical fiber to the Total Internal Reflection Fluorescence (TIRF) illumination arm of the microscope, and laser powers are controlled through an Acousto-Optical Tunable Filter (AOTF) driven by Metamorph. AF647 fluorophores were excited with the 647 nm laser line through a 4-band beam splitter (BS R405/488/561/635, SemROCK). Pumping of AF647 dyes into their triplet state was performed for several seconds using ∼60 mW of the 647 nm laser at the objective front lens. Then, a lower power (∼20 mW) was applied to detect the stochastic emission of single-molecule fluorescence, which was collected using a 100X/1.49 N.A. oil immersion objective, a FF01-676/29 nm emission filter (SemROCK), and an EMCCD camera (Evolve, Roper Scientific). Ten streams of 4,000 frames each on the center quadrant of the camera (256 x 256 pixels of 160 nm side) were acquired at 50 Hz (20 ms exposure time) using Metamorph, representing a total time of 800 s = 13.3 min, during which the 405 nm laser power was progressively increased by a few percent (< 200 µW) so as to keep a relatively constant density of single molecules that emit fluorescence over time. Analysis of the image stacks was made offline under Metamorph, using the PALMTracer program based on wavelet segmentation for single-molecule localization ^65^. Unique super-resolved images of 32 nm pixel size (zoom 5 compared to the original images) were reconstructed by summing the intensities of all localized single molecules (1 detection per frame is coded by an intensity value of 1). The localization precision of our imaging system in dSTORM conditions is around 60 nm (FWHM).

### Photobleaching steps of immobilized GBP*-AF647

GBP*-AF647 from a 7.1 µM stock solution was diluted 1 to 10^5^ in 1 mL PBS, centrifuged to 14’000 rpm for 10 min to remove potential protein aggregates, and the supernatant was placed for 5 min at room temperature in a Ludin chamber containing a clean 18 mm coverslip. After rinsing 2 times with PBS, immobilized proteins were excited with the 647 nm laser under TIRF illumination on the same microscope described above for dSTORM (power 3 mW at the objective front lens). Acquisition sequences of 600 frames each were recorded at 33 Hz in different regions of the coverslip (total time 18 s). Laser illumination caused the photobleaching of all immobilized GBP*-AF647 molecules in this time range. Images were analyzed using Metamorph by measuring the average fluorescence intensity in small regions drawn around individual molecules, and the number of photobleaching steps was determined visually for a total of 276 molecules, on a total of 14 fields of view.

### Primary mouse neuronal cultures and immunoelectron microscopy with GBP*-gold

For the mixed neuronal cell culture from GFP-βNrxn KI mice, pups were sacrificed by decapitation at postnatal day P1-P2. Briefly, hippocampi were dissected and trypsinized (0.25 % trypsin). After the addition of DNase I, cells were suspended in plating medium (DMEM, 10% FCS 1% Glutamate). Cells were plated on Aclar foils coated with poly-L-lysine at a density of 23,333 cells/aclar foil. After 1 h of plating at 37°C, Aclar foils were transferred into neuronal medium (Neurobasal A, 2% GlutaMAX, 2% B27, 0.1 M NaPyr). Cultures were maintained at 37°C in a humidified incubator with an atmosphere of 95% air and 5% CO_2_. Experiments were performed at DIV18 by incubating cultures with GBP*-gold nanobodies for 10 min at 37°C, followed by washing (0.1 M PB), and fixation with 0.1% glutaraldehyde (Serva)/2% paraformaldehyde (Merck) in 0.1% PB. The nanometer gold was subsequently silver-enhanced using the LI Silver Kit (Nanoprobes) for 12 min at RT, followed by an embedding procedure for TEM. After contrasting with osmium (1%), neurons were rinsed with dH2O and dehydrated in an increasing series of ethanol. Cells were incubated with propylene oxide (Electron Microscopy Science) for 15 min, infiltrated with propylene oxide/ epon (1:1) for 30 min, in pure epon resin (EMS) for 3 hours, and hardened at 60 C for 24 h. Ultrathin sections (70 nm) were contrasted with uranyl acetate and lead citrate the following day. Samples were investigated with a transmission electron microscope (TEM, Libra 120, Zeiss) at 80 kV, and images taken with a 2048 x 2048 CCD camera (Tröndle), and with a Hitachi H7650 TEM equipped with an Orius SC1000 (Gatan Inc.) CCD camera.

### Culture of hippocampal neurospheres from H11-Cas9 mice

Hippocampal neurons from H11-Cas9 P0 mice were prepared as previously described ^64^, with modifications. Briefly, pups were anesthetized on ice and sacrificed by decapitation. Dissected hippocampi were treated with papain for 10 min at 37°C, then DNAse for 5 min, before being dissociated in Hibernate-A medium (Gibco, A1247501). The suspension was plated at a density of 500 cells per well, in a U-bottom Ultra Low Attachement 96-well plate (Nunclon Sphera, ThermoScientific 174925), in Neurobasal A medium supplemented with 1.5% heat-inactivated horse serum, 0.5 mM GlutaMAX and B-27 Plus (Gibco, 26050088, 12349015, 35050061, A3582801). This procedure results within 1 day in the formation of compact spheroids containing a mix of neurons and glial cells, of highly reproducible dimensions (150 µm in diameter). To edit the *Gria2* gene coding for the GluA2 subunit, we used the pAAV-SEP-GluA2-TKIT plasmid ^41^, a gift from R. Huganir (Johns Hopkins, Baltimore, MD, USA) (Addgene plasmid # 169442). This plasmid was packaged into an AAV1-U6dblgRNA-SEP-GluA2TKIT produced by the IMN virus production facility from the Bordeaux Neurocampus at a concentration of 1.4x10^14^ gcp/mL. Neurospheres were infected at DIV3 with those AAV1s diluted at 30,000 MOI in culture medium, then maintained in a 37°C - 5% CO_2_ incubator for 2 weeks.

### Labeling and imaging of hippocampal neurospheres from H11-Cas9 mice

At DIV14, neurospheres were collected in a 15 mL tube filled with 2.5 mL warm culture medium and centrifuged at 300 rpm for 3 min in a horizontal swinging bucket centrifuge. Half of the neurospheres were resuspended in 200 µL Tyrode solution supplemented with 1% BSA and containing GBP*-gold at a 1:50 dilution, and left at 37°C for 20 min. Then, 10 mL of warm Tyrode was added, neurospheres were centrifuged again, and this procedure was repeated once. Finally, neurospheres were fixed in 4% PFA-sucrose and 0.2% glutaraldehyde in PBS for 24 h, then processed for TEM. The other half of the neurospheres, fixed without previous labelling by GBP*-gold, was collected under a stereomicroscope and mounted between a microscope slide and a coverslip in Rapid Clear 1.52 solution (Sunjin Lab RC152001), using a 150 µm-thick circular iSpacer (Sunjin Lab, IS006). The SEP-GluA2 signal emitted by infected neurons was observed using an upright confocal microscope (Leica SP-5) equipped with a 60x/1.40 NA oil immersion objective, a 488-nm laser and a photodetector with collecting window between 500-550 nm. Image formats of 2048 x 2048 pixels with zoom factor of 1.5 were selected. Image stacks were acquired with an accumulation of 2 lines at a 400 Hz scanning speed and a pinhole set to one Airy disk, over half of the neurosphere height using a step size of 0.3 µm. Maximal intensity projections were made under FIJI.

### Transmission electron microscopy

Cells on coverslips, or neurospheres collected by sedimentation, were subjected to silver enhancement for 10 min (LI Silver, Nanoprobes Inc.) in the dark. Samples were post-fixed with 1.5% OsO₄ and 1.5% potassium ferrocyanide for 30 min on ice. Sequential dehydration was carried out in 70%, 90%, 95%, and 100% ethanol, followed by one incubation in 100% acetone. Samples were then infiltrated for 2 h with a 1:1 mixture of acetone and Epon resin (Embed-812, EMS), followed by 100% Epon resin for 24 h, and cured at 60 °C for 48 h. Ultrathin sections (70 nm) were cut with an ultramicrotome (Leica EM, UC7) and collected on EM grids. TEM imaging was performed in high-contrast mode using a Hitachi H7650 transmission electron microscope equipped with a CCD camera (Orius SC1000, Gatan Inc.). The number of silver particles per synapse was manually quantified in ImageJ by applying a common detection threshold across all images after background subtraction.

## Acknowledgements

The authors P. Scheiffele and R. Huganir for the generous gift of plasmids; the IINS Cell Biology Facility, especially E. Verdier, A. Caralp, L. Leroy, and N. Retailleau, for primary neuron cultures; R. Sterling, A. Castets, and Z. Andrieux for logistics and cell lines; C. Desquines, B. Tessier, N. Chevrier, and C. Breillat (IINS) for molecular biology; B. Chauvineau for help in neurosphere mounting; J.B. Sibarita, A. Kechkar, and C. Butler for the gift of the PALM-Tracer and WAVE-Tracer programs; P. Costet, C. Martin, and H. El Oussini at the Plateforme In Vivo Exempte d’Organisme Pathogène Spécifique of the Bordeaux Neurocampus (PIV-EOPS) for maintenance of the mouse strains; M. Petrel, S. Lacomme, and M. Fernandez-Monreal for help in electron microscopy.

This research was supported by grants from the French Agence Nationale de la Recherche (ANR-20-CE11-0006-01 “NanoSynAtlas” and ANR-21-CE11-0019-01 “Synaptoligation” to O.T., ANR-21-CE44-0013 “AFFLIGEM” and ANR-23-CE11-0007 “ENDOGRAAL” to M.S.), the GPR LIGHT (post-doctoral fellowship “016-CLEM-probe-2022”) from the University of Bordeaux (Idex program), and a Marie Skłodowska-Curie individual fellowship (101106943) to C.D., and by a grant from the Deutsche Forschungsgemeinschaft (DFG MI 479/10-1) to M.M. Confocal microscopy and part of the electron microscopy were done at the Bordeaux Imaging Center, a node of the FranceBioImaging national infrastructure (grant ANR-10-INBS-04-0). The Cell biology facility of our Institute is funded by grants from the University of Bordeaux (GPR BRAIN_2030).

## Ethical statement

The authors declare that they have complied with all relevant ethical regulations (study protocol approved by the Ethical Committee of Bordeaux CE50).

## Supplementary figure legends

**Figure S1.**
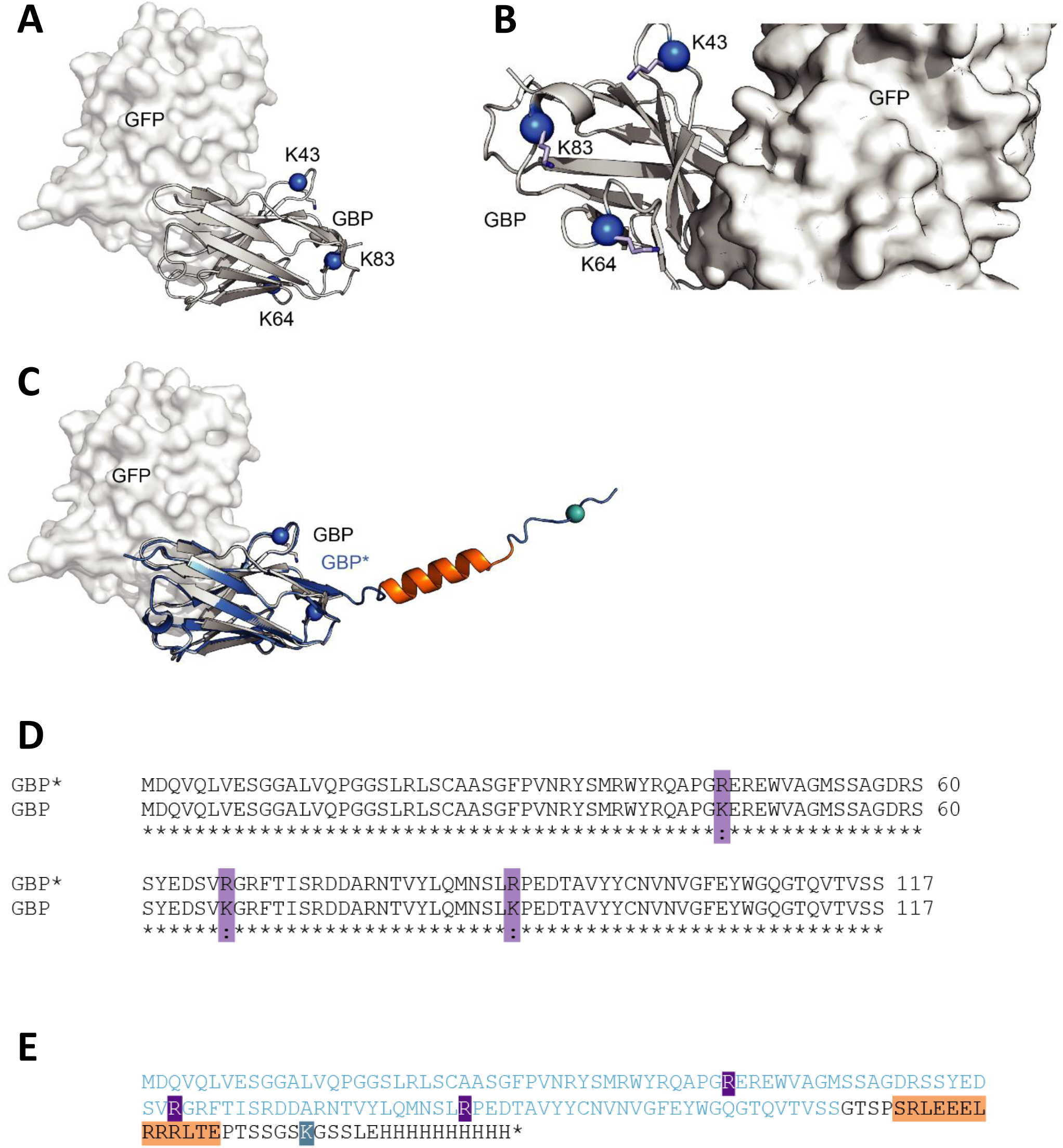
GBP* design. **(A)** Structure of the GFP/GBP complex (PDB: 3K1K). Surface representation for GFP and cartoon for GBP. Positions of three lysine groups (K43, K64 and K83) on the nanobody are highlighted with a blue ball representation for the corresponding alpha-carbon and stick for the side chains. **(B)** Zoom on the GBP/GFP interface. The three lysines on GBP are not directly involved in the GFP binding. **(C)** Overlay of the GBP/GFP complex (white/grey shades) with GBP* alphafold3 structure prediction (blue/orange shades). **(D)** GBP and GBP* sequence alignment. Mutated sites (K to R) are highlighted in purple. **(E)** Sequence of the expressed GBP*. Same color-code as in C. GBP* is in blue with mutated R in purple, alfa-tag is in orange and single lysine group in teal.

**Figure S2.**
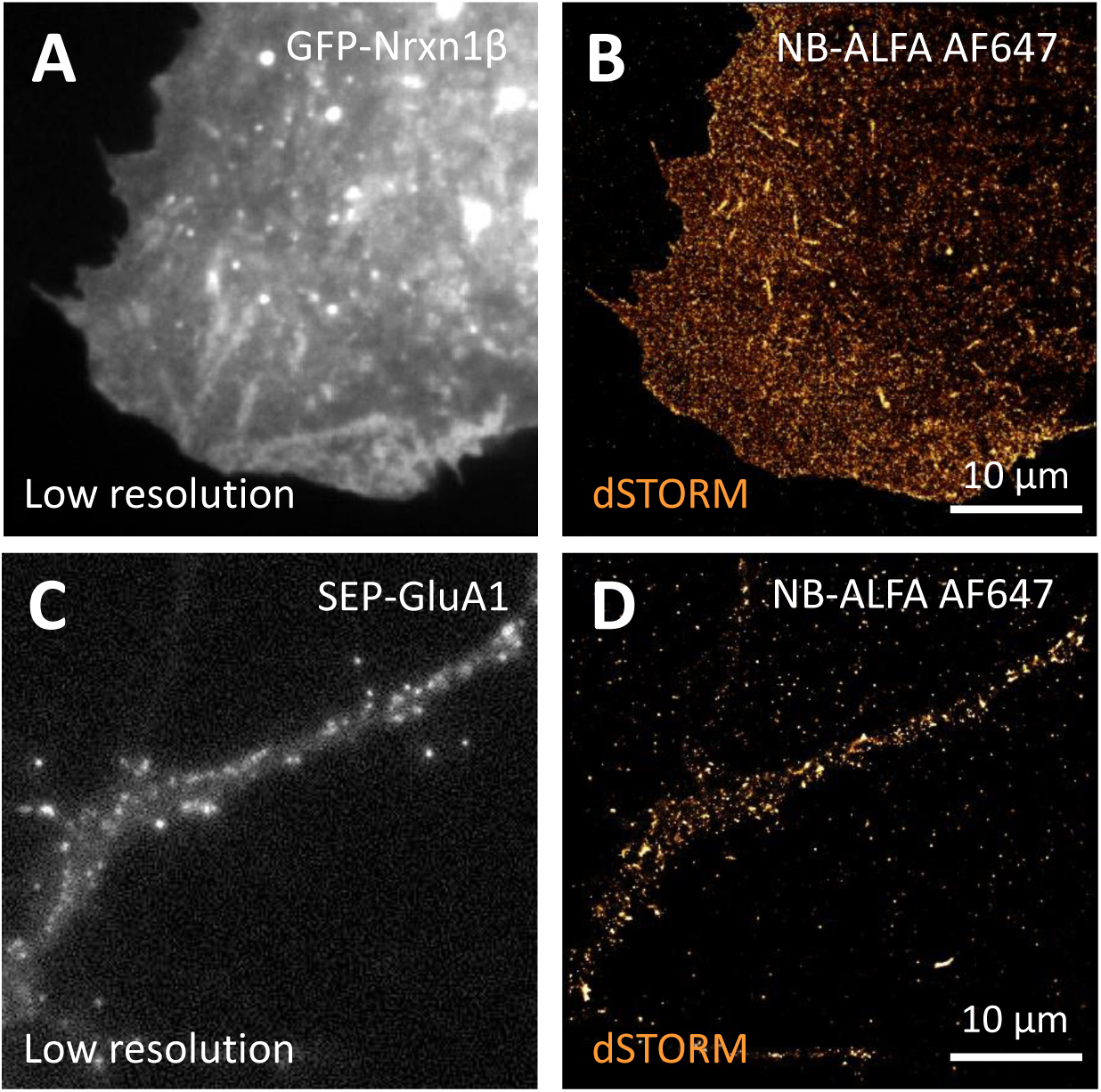
Use of the Nb-ALFA-AF647 to perform dSTORM experiments. **(A,C)** Low-resolution epifluorescence images of a COS-7 cell expressing GFP-Nrxn1β, and of a rat hippocampal neuron expressing SEP-GluA1, respectively. In both cases, cultures were sequentially stained with GBP*-gold, then with Nb-ALFA-AF647, and dSTORM acquisitions were performed on the stochastically emitting AF647 dye (5 rounds of 4,000 images acquired in stream mode at 50 Hz). **(B, D)** Corresponding reconstructions of all localizations obtained from individual Nb-ALFA-AF647 molecules. The zoom factor is x5, i.e. the pixel size is 32 nm compared to 160 nm in panels (A, C).

**Figure S3.**
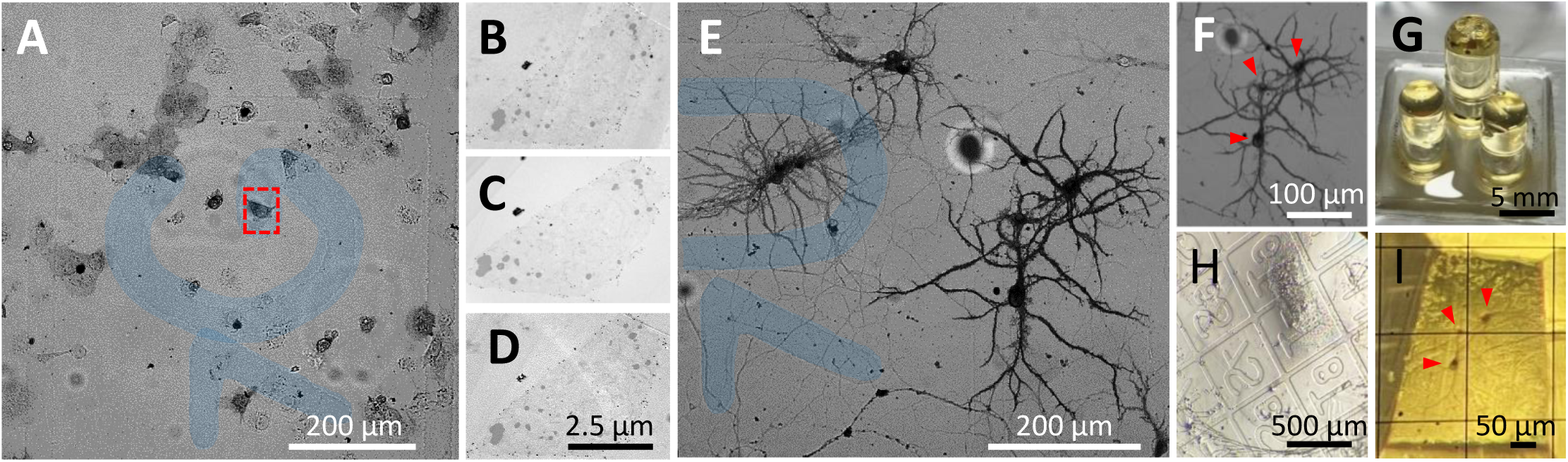
Workflow for correlative light and electron microscopy using etched coverslips. **(A)** COS-7 cells were cultured on etched glass coverslips, which generate precise fiduciary landmarks that enable relocation of the same cells across light and electron microscopy. This correlative approach provides a robust and reproducible workflow to bridge live-cell or fluorescence imaging with ultrastructural analysis, ensuring precise identification of the same cells across modalities. **(B-D)** TEM analysis of a COS-7 cell relocated using the etched landmarks, the three serial ultrathin sections of 70 nm enabling high-resolution ultrastructural visualization. **(E, F)** The same strategy was applied to neurons, demonstrating the versatility of the method. **(G-I)** After resin embedding, the resin penetrated the etched grooves of the coverslip, providing permanent coordinates that guided trimming and ultramicrotome sectioning.

**Figure S4.**
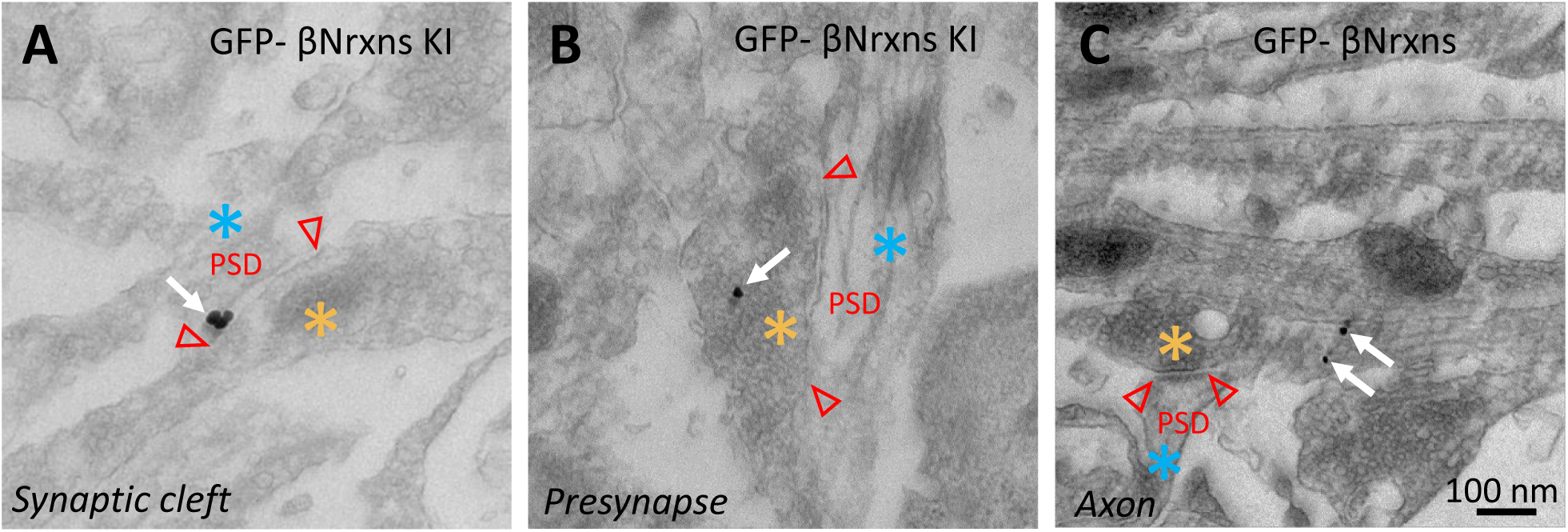
Ultrastructural localization of endogenous β-Nrxns using GBP*-gold. Sample images of neurons from KI mice expressing N-terminally GFP-tagged, endogenous Nrxn1β and Nrxn3β, and labeled with silver-enhanced GBP*-gold. **(A-C)** Detection of β-Nrxns at the synaptic cleft (A), within a presynaptic terminal containing synaptic vesicles (B), and along an axonal shaft (C). Silver-enhanced gold particles are shown with arrows.

## Notes

### Competing Interest Statement

The authors have declared no competing interest.

